# Are ants always mutualistic? Influence of ants and variation in flowerets of *Clerodendrum chinense*

**DOI:** 10.1101/2020.02.29.971457

**Authors:** Rupesh Gawde, Nivedita Ghayal

**Author notes:** Corresponding Author: NIVEDITA GHAYAL, Department of Botany, Abasaheb Garware College, Pune, India.

## Abstract

Ants are the major mutualists in the plant -ant interaction system playing a major role in the survival of plants by deterring herbivore and increasing plant health and fitness of the plant. To get better insights into the ant-plant interaction system, *Clerodendrum chinense* occupied by *Crematogaster roghenoferi* were observed for six months to assess the influence of presence and absence of ants on the flowerets blooming. In a total of (n=34) flowerets bunch, 17 were used as control and 17 experimentation that were treated with ant barrier glue to restrict the ants visitation to the flowerets bunch. By using the Students t-test, our study revealed the facultative behaviour of ants. Citing the equal presence of herbivores in both the control and experimented sets, and decreased floweret blooming in anting-continued (control) whereas increased floweret blooming in de-anted (experimented). The present study has shown the facultative interaction of *C. roghenoferi* with *C. chinense* and its effects on the reproductive structures (flowerets) of the shrub. This study makes a clear case stating that not always an interaction between ant and plant is the mutualism.

## Introduction

Mutualism is commonly defined as an interaction between two unrelated species that positively affects the fitness of both species, where, the fitness of a population is defined as the success of a population in disseminating its own genes (Jorgensen, 2008). An example of a mutualism that directly influences reproductive success is the relationship between plant and pollinator (Bascompte, 2019), Where a pollinator benefits the plant in its reproductive success. In turn, pollinators gain resources, either by gathering nectar or by exploiting a potential plant to lay eggs on, thereby increasing its reproductive success as well (Nepi et al, 2018). In interspecific species interactions, the amount of mutualist dependence on each other ranges from obligate to facultative (Mukhopadhyay & Quader, 2018; Silknetter et al, 2020). Facultative mutualism is when two species live together by choice, whereas, obligate mutualism is when two organisms are in a symbiotic relationship because they cannot survive without each other (Liu et al, 2010). Ant and Plant mutualisms are of great interest in the ecological research field, Ants are social insects dominating most terrestrial habitats (Sudd & Franks, 1987). They forage at every level of the forest, humus, and soil and in many cases show a mutualistic relationship (Bronstein et al., 1998). Ant-plant associations have arisen independently in many plant taxa and often thought to represent good examples of coevolution, even if not, they provide dramatic examples of species mutualism and adaptation (Bronstein et al., 2006). The extrafloral nectary (EFN) is regarded as a general and reward capable of attracting to the plant ants with wide diversity (Beattie, 1985). Some plants attract ants via extrafloral nectaries that act as sources of sugars for protection against insect herbivores (Grasso et al., 2015). A variety of nectar-feeding insects are attracted by EFNs (Marazzi et al., 2013), which also attracts Insect herbivores that plays an important role in several factors of host plant fitness (Jones et al., 2017). Ants are by far the most common visitors to EFN-bearing plants both in temperate and tropical habitats (Agarwal & Rastogi, 2008). Such plant species are termed as myrmecophyte providing dwellings that are, domatia, for ants in the form of swollen hollow internodes, thorns, stipules, leaf pouches or chambers within epiphytic tubers (Gaume et al., 2005). Ants are the major exploiters of such domatia collecting food bodies, from modified trichomes also sometimes called extrafloral nectaries (EFNs) with high lipid and protein contents that are specifically adapted to meet the ants’ nutritional requirements (Chanam et al., 2014). Extrafloral nectaries (EFNs) are nectar secreting organs not directly involved in pollination which are found virtually only above ground-plant parts (Mizell RF & Mizell PA 2008). In the past, many experimental field studies have demonstrated that ant visitation to EFNs may increase plant fitness by deterring bud or flower herbivores (Rico-Gray & Thien 1989; Oliveira 1997; Del-Claro et al., 2016; Do Nascimento & Del-Claro, 2010). Ants can be very efficient in reducing insect herbivory in several plant species that in turn increases the fitness level of the said plant species (Do Nascimento & Del-Claro, 2010, Bentley et al., 1977). However, these obligatory interactions between plants and EFNs may be facultative as argued by (Speight et al 1999). To get more insights into the presence of such facultative interactions, in our present study we examined and observed the interactions between a perennial shrub *Clerodendrum chinense* and, an arboreal ant species *Crematogaster roghenoferi* to see if there is the presence of mutualism i.e. Do the presence of ants reduce herbivores and increase the number of blooming flowerets?

## Material and methods

### Study area

Fieldwork was carried out between June-August 2018 and January-February 2019 in the flowering season, at a Biodiversity Park (19.19979 N; 72.86169 E) in Mumbai, Maharashtra. Climate is mostly warm and humid, the mean annual temperature ranges from 30-32°C.

### Methodology

Ramets were observed from bud growth to floweret maturation. In 32 visits, (n = 34 ramets) 17 were sampled for (experimentation) de-anted (DA) and 17 for (control) anting-continued (AC) to the field site. All biotic aspects like water, temperature, etc. for both the sets were kept constant.

Identification for the shrub was done using the key from (Leeratiwong et al., 2011) and ant species were identified using the (Hölldobler & Wilson, 1990) key. Herbivory presence in the flowerets was scanned visually for a fixed time range and insect herbivores were collected and preserved in 70% ethanol. Moreover, identified until order level (Schell & Latchininsky, 2007) (insects were considered to be herbivores if they were found to be feeding/damaging the plant).

The de-anting procedure was carried out with a sticky barrier made with a mixture of Vaseline (Unilever products) and peppermint oil 10 ml (Dr. Jain’s forest herbal Pvt. Ltd) that had a success rate of 11 days. (Success rate was calculated on the basis of an ant crossing the barrier in days after application of barrier). Then applied on the stems over a thin plastic film of ramets to stop the intrusion of ants towards experimented (De-anted flowerets). The bloomed and withered flowerets were counted from control and experimented bunch to assess the difference and herbivory effect. All statistical analysis were performed in Microsoft excel 2013.

## Results

### Shrub Identification

The shrub species were identified as *C. chinense* a perennial introduced shrub native to China, a major weed found on roadsides, gardens, towns, and villages commonly used as an ornamental plant for its sweet-smelling inflorescence.

### Ant identification

The species of ant found on the shrub was identified as *C. rogenhoferi* Mayr, 1879 using the key (Hölldobler & Wilson, 1990) an arboreal species very common to the Indian sub-continent. They forage for several types of plant-derived food resources, including floral and extrafloral nectar from many plant species.

### Maturation difference

There was no significant difference between the Maturation period of the De-anting and Anting-continued ramets (*P*-value=0.08). The effect of the de-anting procedure (application of ant barrier glue) did not show any significant difference, stating that the newly made ant barrier glue did not affect the maturation period (Fig. 1).

**Figure. 1.**
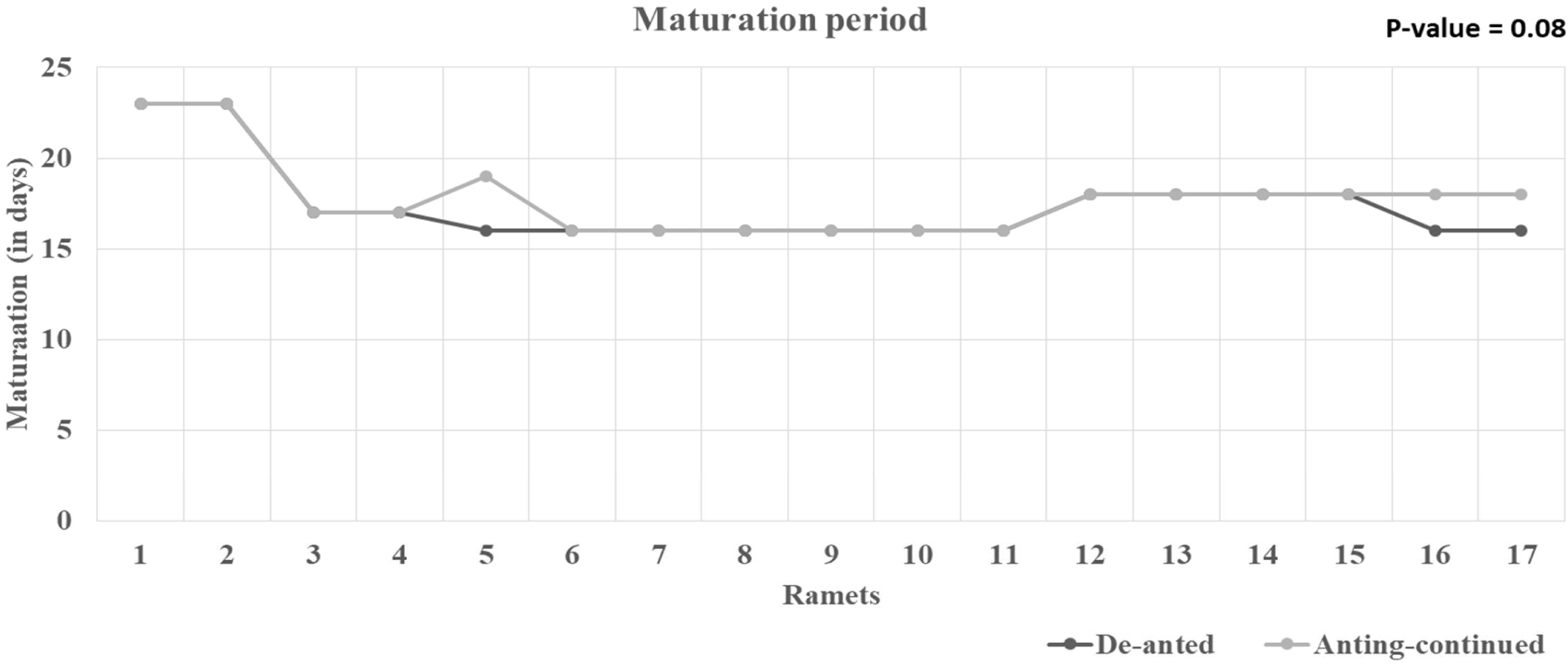

### Budding difference

Budding was significantly higher in the experimented (de-anted) bunch and less in control (anting-continued) bunch. The student t-test showed a significant difference between total budding of De-anted and an anting-continued bunch (*P*-value = 0.003). De-Anted bunch, showed the highest budding in the second quartile at 38 buds and lowest in the first quartile at 14 buds, with a median at 26 and mean of 26.35. Anting-continued bunch showed the highest budding in the second quartile at 28 buds and lowest in the first quartile with the lowest value at six buds, with a median at 16.5 and mean of 17.2 (Fig. 2). The significant difference between total budding in de-anting and Anting-continued ramets shows the increased budding in the absence of ants and decreased budding in the presence of ants.

**Figure. 2.**
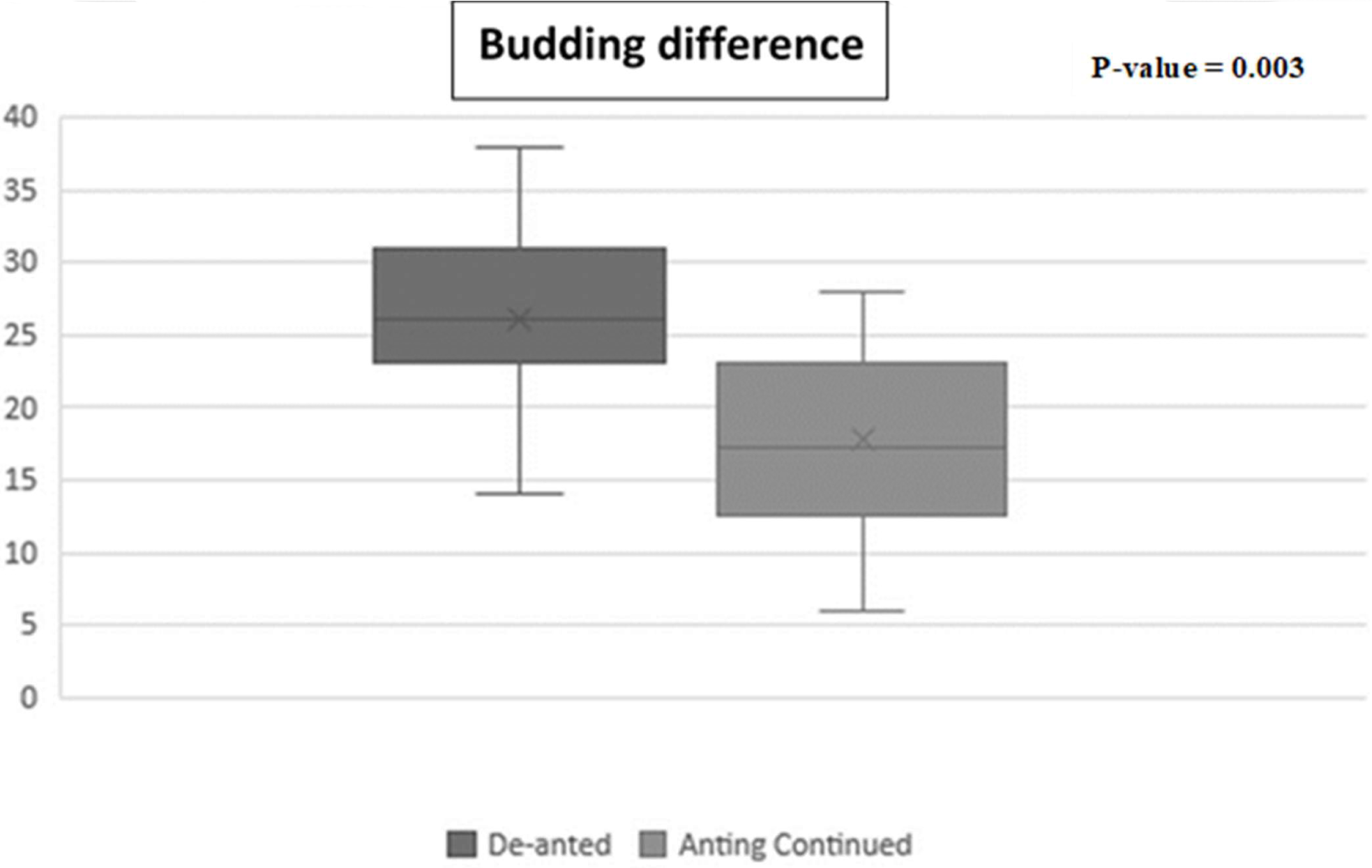

### Blooming difference

Blooming was significantly higher in the experimented (de-anted) bunch and less in control (anting-continued) bunch. The student t-test showed a significant difference between total blooming of De-anted and an anting-continued bunch (*P*-value = 0.002). De-Anted bunch showed the highest budding in the second quartile at 22 bloomed flowerets and lowest in the first quartile at eight bloomed flowerets, with a median at 17 and mean of 16. 23. Anting-continued bunch showed highest blooming in the second quartile at 20 bloomed flowerets and lowest in the first quartile with the lowest value at zero bloomed flowerets, with a median at 10 and mean of 9.82 (Fig. 3). The significant difference between total blooming in de-anting and Anting-continued ramets shows the increase in floweret blooming in the absence of ants and a decrease in floweret blooming in the presence of ants

**Figure. 3.**
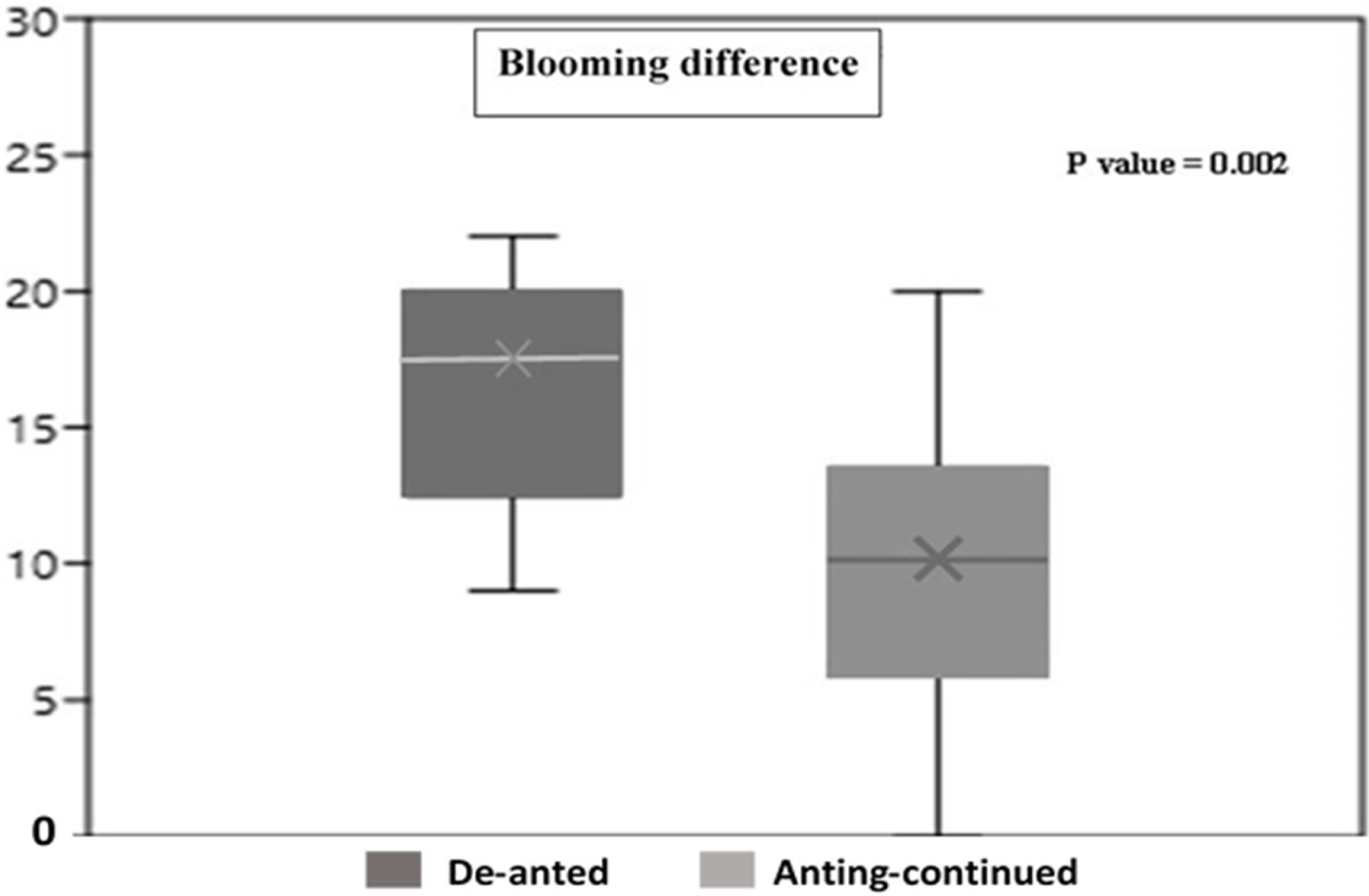

### Herbivore diversity

A total of 13 insect herbivores were identified to their order level out of which (in percentage) Hemipterans (31%), and Coleopterans (31%), Orthopteran (7%), Lepidopterans (23%), and Bllatodea were (8%) The herbivore diversity was assessed by using (Shannon Diversity index = 1.278) (Fig. 4). The student t-test for experimented (de-anted) and control (anting-continued) herbivores showed no significant difference between herbivore data set (*P*-value = 0.61) (Fig. 5). No significant difference in the number of herbivores was found on both de-anting and Anting-continued ramets, stating that the presence of ants and absence of ants did not have any effect on herbivores. Hence confirming the presence of facultative interactions between *C. chinense* and *C. roghenoferi*

**Figure. 4.**
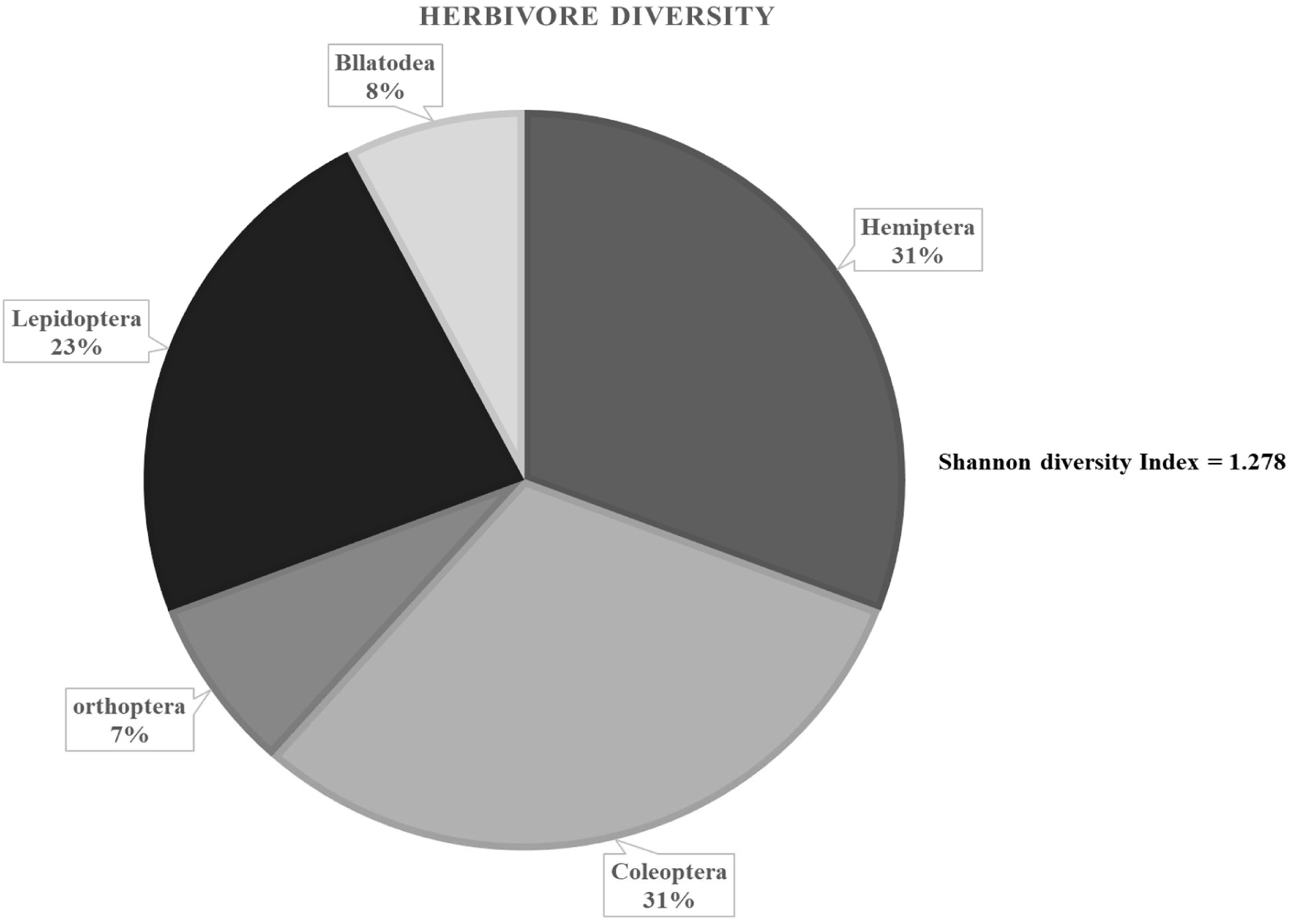

**Figure. 5.**
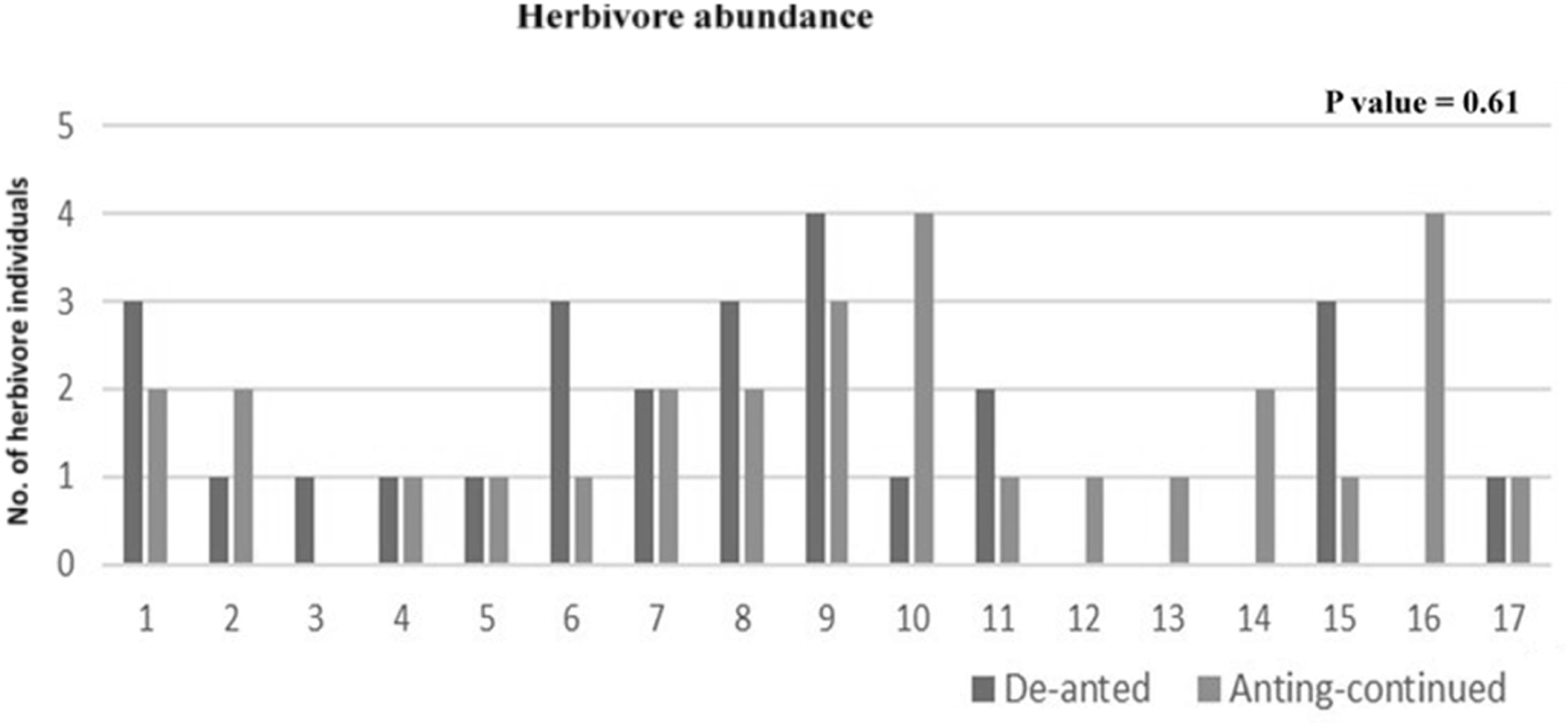

## Discussion

EFN visitation of ants have been reported in the botanical literature. Extrafloral nectar may signify an extremely rich food source for ants because it contains sugars (fructose, glucose, and sucrose) and several amino acids, which are essential for insect nutrition. The current study involving *C. chinense* and *C. roghenoferi* and insect herbivores extends earlier studies by providing a more detailed picture of the species intersections within the system through de-anting experiments. The extrafloral nectaries are regarded as a reward capable of attracting plants with a varied diversity of ants (Beattie et al, 1985), some studies also showed the round the clock species turnover of ant species visitation to EFNs (Oliveira et al., 1999). However, the present study showed the presence and dominance of only *C. roghenoferi*, throughout the study period. Studies have shown the capacity of EFNs to reduce the herbivory by insects by attracting mutualistic ants. This is the first demonstration of ants displaying facultative interaction with an EFN bearing shrub *C. chinense*, where ant visitation leads to decreased flowerets under natural conditions.

Foraging ants are known to attack and dislodge bud/fruit feeding hemipterans on plant bearing EFNs near reproductive structures (Ness, 2006). Similarly, *Crematogaster opuntiae* ants are known to attack/kill Chelinidea *sp* and sucking bugs on *O. acanthcarpa* (Pickett & Clark 1979). These reports directly contradicts our results where, equal number of herbivores were found to be present on experimented (de-anted) and control (anting-continued) flowerets containing EFNs. Our de-anting experiments showed the equal presence of herbivores in both (de-anted) and control (anting-continued) flowerets, this result directly correlates and supports a study that showed herbivores possessing mechanisms to overcome ant predation and can feed on the plant despite the presence of ants (Horvitz & Schemske 1984). Our study also supported the study demonstrating variation in the abundance, aggressiveness or size of ant visitors can affect their protective abilities against herbivore species (Bentley 1977).

## Conclusion

Although the type of herbivory damage caused by herbivores remains unknown citing the difficulties in measuring standards to assess the direct damage to the plant reproductive structures. The present study has shown the facultative interaction of *C. roghenoferi* with *C chinense* and its effects on the reproductive structures (flowerets) of the shrub. This study makes a clear case stating that not always an interaction between ant and plant is mutualism

## Acknowledgments

The authors are thankful to Principal, MES Abasaheb Garware College, Pune for providing the facilities for carrying out this research work. We would also like to thank Dr. Tejaswani Pachpor for helping in the identification of ant species, Dr. Pravin Tambe for assistance in data analysis. Dr. Dhanashree Paranjpe, Dr. Sumedha Korgaonkar, Dr. Ankur Patwardhan, Ms. Monali Mhaskar for their suggestions to improve the nature of the study. In addition, staff and Colleagues of Annasaheb Kulkarni Department of Biodiversity, MES Abasaheb Garware College, Pune for their constant support.

